# The Heterochromatin protein 1 is a master regulator in RNA splicing precision deficient in ulcerative colitis

**DOI:** 10.1101/2020.12.30.424798

**Authors:** Jorge Mata-Garrido, Yao Xiang, Yunhua Chang-Marchand, Caroline Reisacher, Elisabeth Ageron-ardila, Chiara Guerrera, Inigo Casafont, Aurelia Bruneau, Claire Cherbuy, Xavier Treton, Anne Dumay, Eric Ogier-Denis, Eric Batsche, Mickael Costallat, Gwladys Revêchon, Maria Eriksson, Christian Muchardt, Laurence Arbibe

## Abstract

Defects in RNA splicing have been linked to numerous human disorders, but remain poorly explored in inflammatory bowel disease (IBD). Here, we report that, in the gut epithelium of patients with ulcerative colitis (UC), the expression of the chromatin and alternative splicing regulator HP1γ is strongly reduced. Accordingly, inactivation of the HP1γ gene in the mouse gut triggered several IBD-like traits, including inflammation and dysbiosis. In parallel, we discovered that its loss of function broadly increased splicing noise, reducing requirement for canonical splicing consensus sequences, and favoring the usage of cryptic splice sites at numerous genes with key functions in gut biology. This notably resulted in the production of progerin, a noncanonical toxic splice variant of prelamin A mRNA, responsible for the Hutchinson Gilford Progeria Syndrome (HGPS) of premature aging. Likewise, production of progerin transcript was found to be a signature of colonic cells from UC patients. Thus, our study identifies HP1γ as a regulator of RNA metabolism *in vivo*, providing a unique mechanism linking anti-inflammation and accuracy of RNA splicing in the gut epithelium. HP1 defect may confer a general disturbance in RNA splicing precision to scrutinize in IBD and more generally in accelerating aging diseases.

## Main

Inflammatory bowel disease (IBD), including ulcerative colitis (UC) and Crohn’s disease (CD), are chronic inflammatory gut disorders characterized by an uncontrolled inflammation leading to bowel damage. While susceptibility gene loci have been identified, genetic factors account for only a portion of overall disease variance, indicating a need to better explore gene-environmental factor interactions in the development of the disease^1^. Epigenetics captures environmental stresses and translate them into specific gene expression pattern and little is known on the role played by chromatin deregulations in the pathogenesis of IBD. The Heterochromatin Protein 1 proteins (HP1α, HP1β, and HP1γ in mouse and human) are readers of the H3K9me2/3 histone modifications. They play key roles in formation and maintenance of heterochromatin, thereby participating in transcriptional gene silencing^2^. In addition to their “caretaker” function in heterochromatin, HP1 proteins display RNA binding activity described across the species^3^. In mammals, *in vitro* studies have shown that HP1γ binds intronic repetitive motifs of pre-messenger RNA^4^ and promote co-transcriptional RNA processing as well as alternative splicing^5–6^. These studies suggest a role in RNA metabolism homeostasis that remains unexplored in intestinal disorders, in sharp contrast to neurodegenerative, cardiac or premature aging diseases^7^. In this study, we have delineated essential functions played by HP1γ in the regulation of gut homeostasis with relevance to IBD, and identify an essential safeguarding function on RNA splicing in the epithelium.

### HP1γ inactivation triggers IBD-like traits

Our previously reported observations on the impact of HP1γ on the control of inflammation in the gut in response to bacterial infection^8^ prompted us to examine expression of this protein in the context of chronic inflammation. To that end, we examined an available cohort of colonic biopsies in non-inflamed tissue from UC patients and from healthy individuals undergoing screening colonoscopies (**Figure 1a and Supplementary Data Table 1 for detailed description of the population).** Quantitative immunofluorescence (IF) showed a strongly decreased expression of HP1γ in the colonic epithelium of UC patients, as compared to healthy individuals (control patients) (**Figure 1a-b)**. Compromised HP1γ expression in association with UC was confirmed using the EXCY2 mouse model, which combines immune dysfunction (IL-10 deficiency) and epithelium NADPH oxidase 1 (Nox1) deficiency^9^. At early stage, these mice exhibited a spontaneous chronic colitis that evolved into a colitis-associated dysplasia and adenocarcinomas. Reduced HP1γ expression in the epithelium prevailed at the initial chronic inflammatory stage (1—5 months aged), while at the cancer stage, the expression was recovered, in coherence with the reported increase detection of the Cbx-protein family members in various cancers^10–11^ (**Figure 1c-d**).

**Figure 1:**
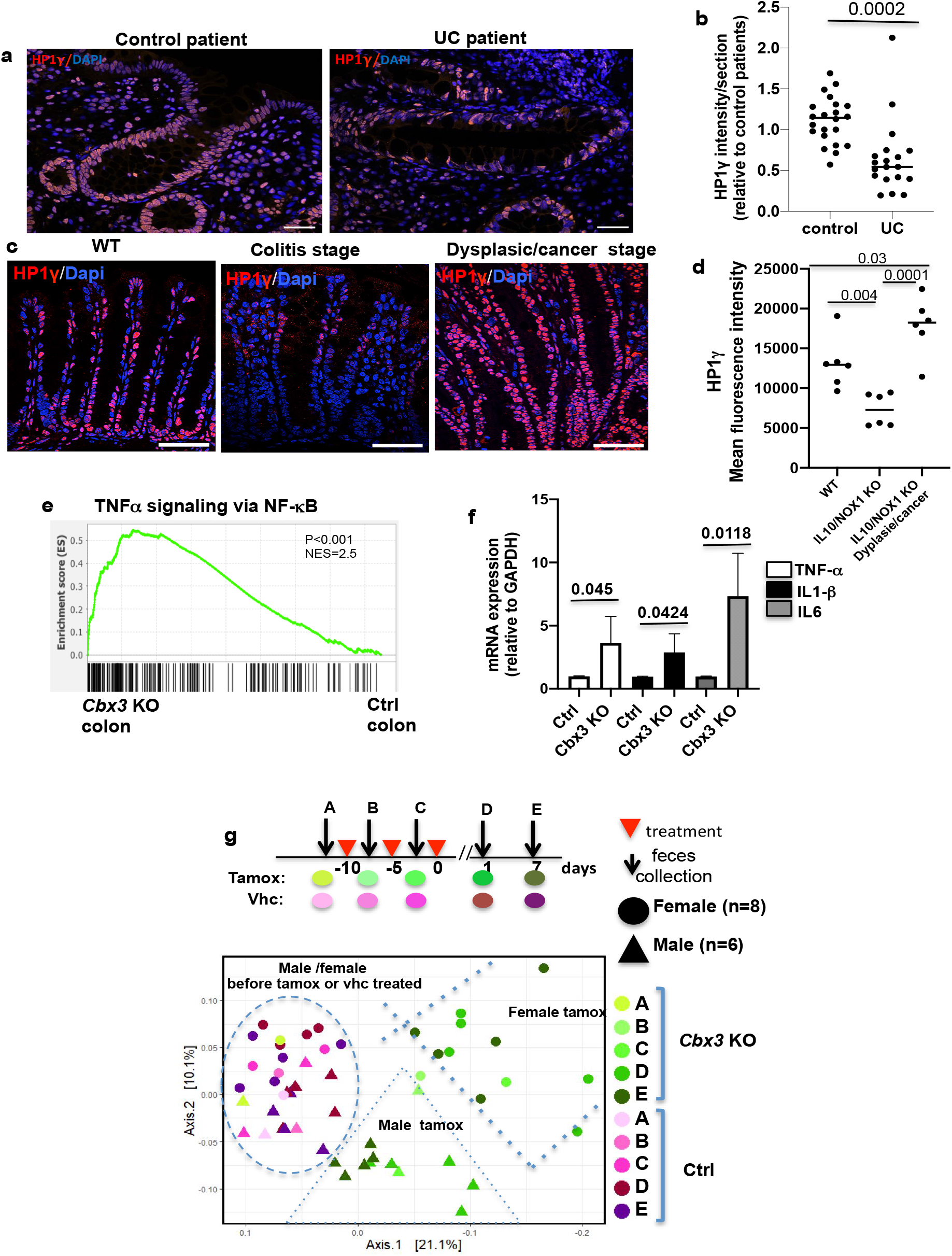
*Cbx3* inactivation in the epithelium leads to gut homeostasis rupture. **(a-b)** HP1γ expression is affected in UC patients: in (**a**) Representative immunofluorescence in colonic tissue sections stained with anti-HP1γ antibody (red) and Dapi (blue) of colon sections (Scale bar: 50μm) (**b**) ImageJ quantification of the mean fluorescence HP1γ signal intensity/section in control (n=10) and UC patients (n=16), expressed as relative value to control patients (Student’s *t* test) (**c-d**) Biphasic expression of HP1γ in the IL10/NOX1 KO mice model: in (**c**) Immunostaining with anti-HP1γ antibody (red) and Dapi (blue) fluorescence, (**d**) ImageJ quantification of the mean florescence Intensity. n=6 mice for each group, Scale bar: 80μm. (Student’s *t* test) (**e)** Significant enrichment of the colon *Cbx3* KO transcriptome with pro-inflammatory signature, Two-sided nominal *P* values were calculated by GSEA. (**f**) mRNA expression of TNF-a, IL1-β and IL6 by RT-qPCR from colon epithelium of Ctrl (control) and *Cbx3* KO mice (n=3-4 mice/group, Student’s *t* test) (**g**) Temporal evolution of the Beta diversity in Villin-Cre *Cbx3* male and female mice. Sheme illustrating the time course of fecal sample collection before (defining group A) and after treatments (Vhc or Tamox) (defining groups B-E), in female (n=8) and male (n=6) mice. Statistical analysis is provided in **Supplementary data Table 3**.

These observations, suggesting a role for HP1γ in chronic inflammation, prompted us to generate a Villin-creERT2:*Cbx3-/-* mouse model (referred to as *Cbx3* KO mice), allowing inducible inactivation of HP1 γ in the epithelial lineage of the gut. In these mice, tamoxifen gavage resulted in complete depletion in the HP1γ protein in the epithelium of the tested tissues, although the depletion was accompanied by an up-regulation of the HP1α and HP1β isoforms (**Extended data Figure 1**), as previously reported^12^. Next-generation RNA-sequencing on purified epithelial cells from either crypts, villi, and colon in young adult mice (8-10 weeks aged) indicated extensive changes in the transcriptome landscape in the *Cbx3* KO mice epithelium, with increased signature scores in pathways involved in lipid oxidation, symptomatic of oxidative damage (**Supplementary Data Table 2)**. Moreover, specifically in the colon, we noted a substantial increase in inflammatory gene expression confirmed by Q-PCR (**Figures 1e-f, Supplementary Data Table 2)**.

**Extended data Figure 1.**
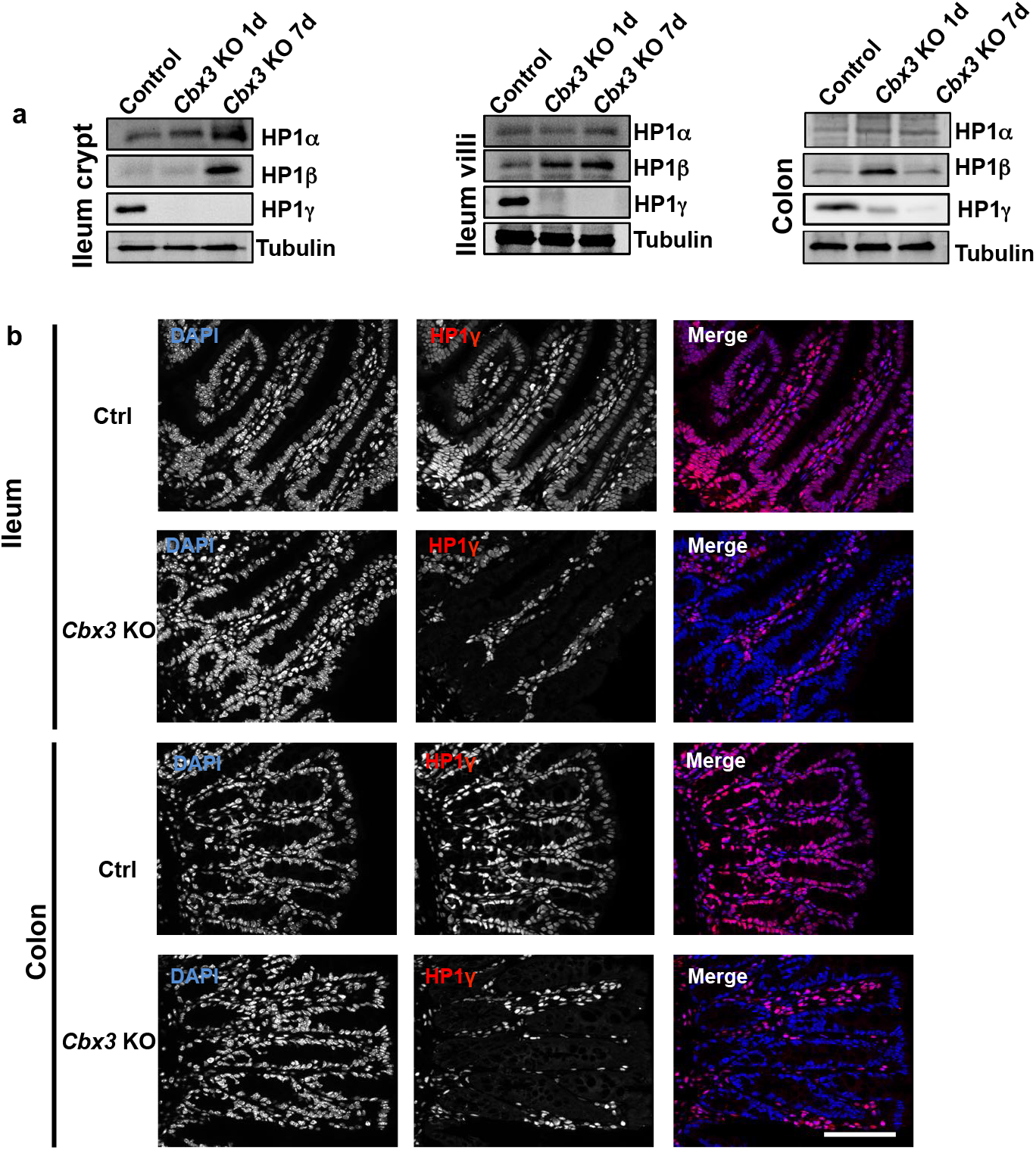
Validation of the *Cbx3* KO mice model. (**a**) Western blot analysis: time course of HP1 isoforms expression upon *Cbx3* Knock down 1 day and 7 day post-tamoxifen gavage in the crypt, villi and colon epithelia (**b**) Immunostaining HP1γ in control (Vehicle treated) and *Cbx3* KO mice showing a loss of HP1γ specifically in the epithelium compartment. Scale bar: 80μm.

As both modified lipid metabolism and inflammation are conducive of gut microbiome dysbiosis^13^, we further characterized the fecal microbiomes in mice (n=6 males and n=8 females) *via* Illumina sequencing of the V3-V4 region of bacterial 16S rRNA. In subsequent UniFrac principal coordinates analysis, *Cbx3* KO mice clustered away from the WTs, indicative of a shift in the microbial communities (**Figure 1g** and **Supplementary data Table 3 for statistics details**). This shift was exacerbated in females (p value=0.006) as compared to the males (p value=0.015), although the bacterial communities were similar in the two sexes prior to *Cbx3* inactivation(**Figure 1g)**. The bacterial composition also remained unchanged in *Cbx3* fl/fl female mice treated with tamoxifen in the absence of the Cre recombinase, ruling out an effect solely induced by tamoxifen administration (**Extended data Figure 2**).

**Extended data Figure 2.**
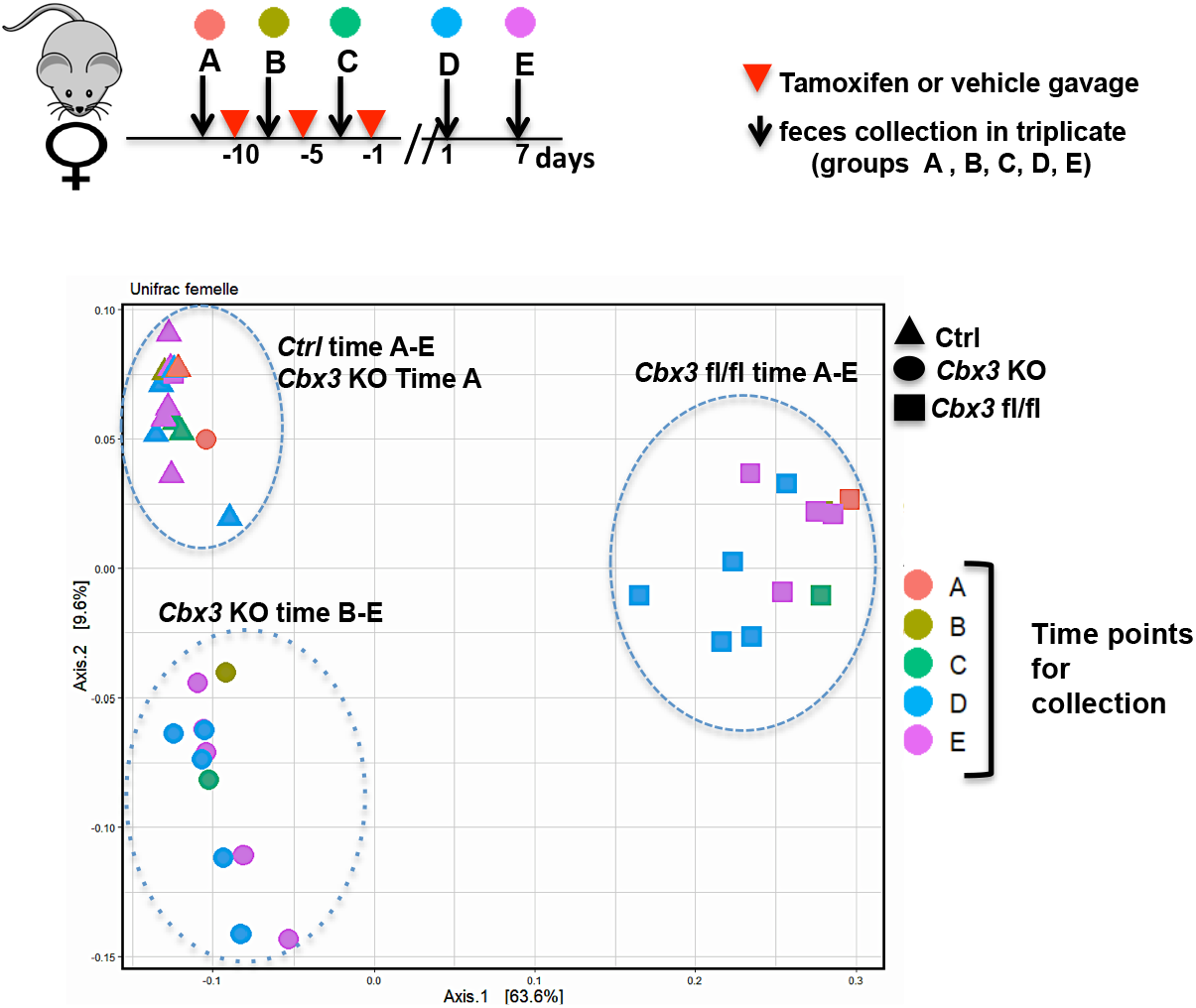
Temporal evolution of the Beta diversity in Villin-Cre *Cbx3* KO female mice before (group A) and after treatments (Vhc or Tamox) (groups B-E): **(a)** Sheme illustrating the fecal sample collection method before and after treatments in the Ctrl (n=4), *Cbx3* KO (n=4)and *Cbx3* fl/fl (n=3) mice (**b**) Beta diversity showing a shift only in the *Cbx3* KO mice. Tamoxifen treatment did not impact beta diversity in the *Cbx3* fl/fl mice that do not expressed the CRE recombinase.

As gender may influence IBD risk factors^14^, we pursued separate analysis of male and female mice. In female mice, 110 OTUs were modulated in response to HP1 inactivation while in males, only 60 OTU covaried significantly with HP1 inactivation (**Supplementary data Table 4**). In females particularly, we observed an overrepresentation of colitogenic bacteria such *as E. coli and Alistipes*, and a profound down-regulation of anti-inflammatory bacterial species such as the short-chain fatty acids (SCFAs) producer *Ruminococcaceae* **(Extended data Figures 3),** both phenomena being symptomatic of a dysbiotic microbiota reported in IBD^15^.

**Extended data Figure 3.**
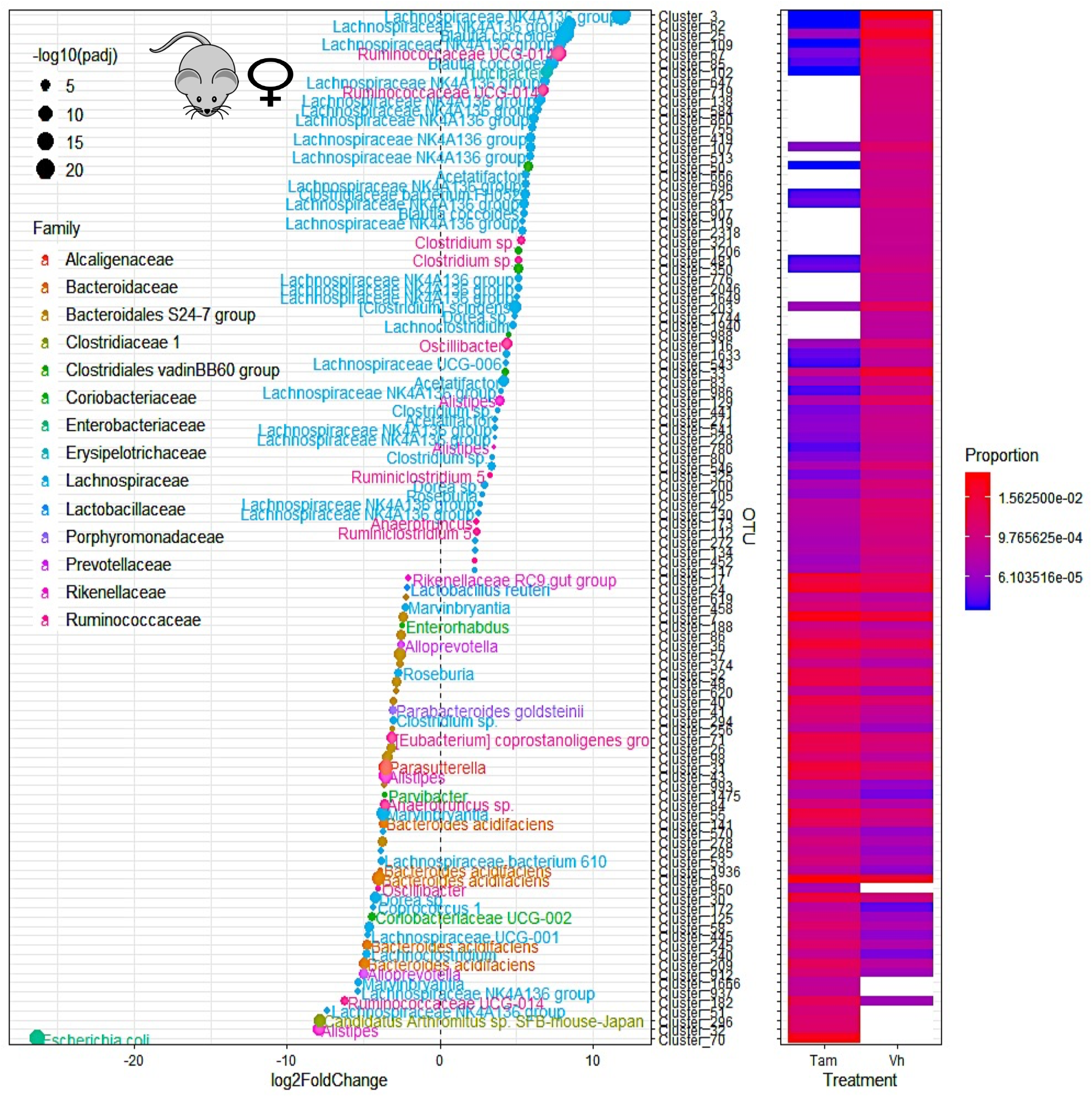
Heat map representing the differentially abundant OTUs between tamoxifen (TAM) and Vehicle (Vh) treated *Cbx3* KO in female mice feces (n=4 in each treated group) with fold change > 2 and significant (P < 0.05). For the graph on the left, each OTU is represented by a dot and colored according to its taxonomic classification at the family level. Taxonomy at the genus or species level is also indicated, when available, next to each OTU. A logarithmic scale (log-2) was used for the x axis

Thus, UC is associated with a reduced expression of HP1γ in the gut epithelium, while inactivation of the cognate gene in the mouse gut results in features typical of gut homeostasis rupture. We concluded that in the colon epithelium, HP1 exerts protective functions, conveying to anti-inflammation and microbiota homeostasis.

### Impact on proliferative homeostasis and maturation

We next delineated the homeostatic functions played by HP1γ in the small intestine. The transcriptome analysis was indicative of a de-silencing of E2F target-genes upon inactivation of *Cbx3* **(Supplementary Data Table 2)**, in agreement with its reported role in retinoblastoma (Rb)-mediated control of cell division^16^. Consistent with an effect of *Cbx3* inactivation on the cell-cycle, the proliferation marker Ki67 in the mutant mice was ectopically detected beyond the normal proliferative compartment, extending all along the crypts and at the base of the villus axes (**Extended data Figure 4a**). Furthermore, a one-hour pulse of the thymine analog 5-bromo-29-deoxyuridine (BrdU), marking cells in S phase, resulted in a frequent labeling of cells at the crypt base containing the intestinal stem cell (ISC) in *Cbx3* KO mice (**Figures 2a-b)**. In WT mice, this labeling was predominantly confined to the immediate ISC progeny compartment *i.e.* the transit amplifying cells, while for the ISC, BrdU was poorly incorporated, consistent with the prolonged G1 phase of stem cells^17^. Expansion of the stem-cell niche in the *Cbx3* KO mice was also documented by an enlarged area of detection for the Olfm4 marker of stemness (**Extended data Figures 4b-c**). Finally, *ex vivo* enteroid 3D matrigel cultures displayed accelerated bud formation when cells were collected from *Cbx3* KO mice (**Extended data Figures 5a-b**). However, upon prolonged culture, the bud per organoid ratio drastically dropped in *Cbx3* KO-derived organoids, possibly as a consequence of cellular exhaustion (**Extended data Figures 5a-b**). In the organoids, detection of cells in S-phase by Edu incorporation also revealed aberrantly positive cells along the villus axes, with EdU positive cells filling the lumen of the organoids. (**Figure 2c and Extended data Figure 5c**).

**Figure 2:**
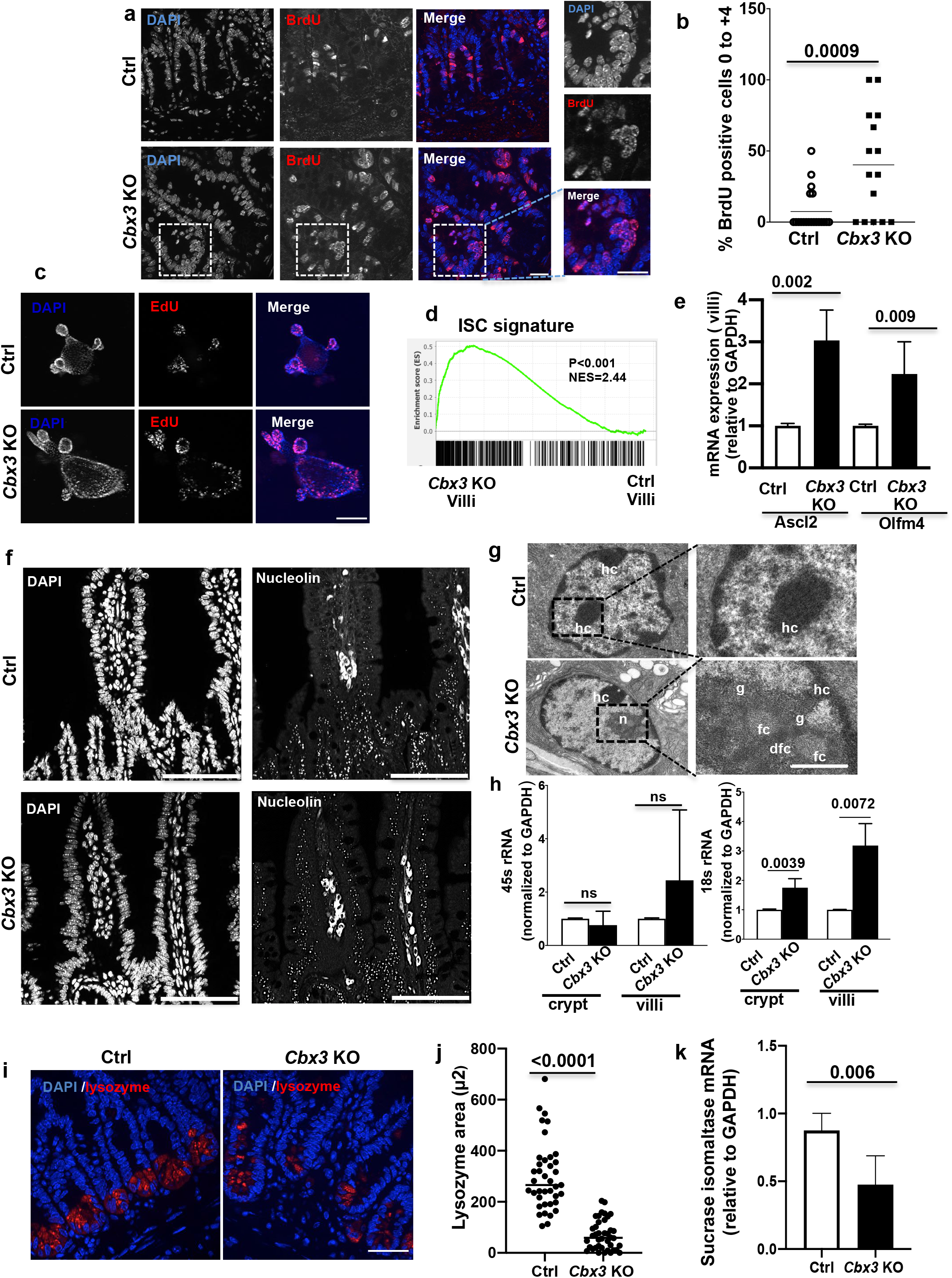
HP1γ controls epithelial proliferation and maturation in the small intestine. **(a)** Immunostaining with anti-BrdU antibody (red) and Dapi (blue) of crypt ileal sections from Ctrl and *Cbx3* KO mice (Scale bar: 20μm) (**b**) Quantification of the BrDU positive cells (red) versus Dapi (blue) at the stem cell compartment (0 to +4 position) n=3 animals Ctrl, n=20 sections; *Cbx3* KO n=15 sections; Student’s *t* test) (**c**) co-staining EdU (red)/Dapi (blue) in organoids derived from Ctrl and *Cbx3* KO mice (Scale bar: 100μm) (**d**) Significant enrichment of the villi *Cbx3* KO transcriptome with the Lgr5+ intestinal stem cell signature (Munoz et al, 2012), Two-sided nominal *P* values were calculated by GSEA. (**e**) mRNA expression of the stemness markers Olfm4 and Ascl2 by RT-qPCR from villi epithelium of Ctrl and *Cbx3* KO mice (n=3-4 mice/group, Student’s *t* test) **(f)** Representative immunofluorescence with nucleolin antibody, n=6 mice/group (Scale bar: 150μm) (**g**) Transmission electron microscopy (TEM) characterizing the nucleolar structure at the upper part of the villi. **g-Left:** Heterochromatin (hc) was observed in the nucleus of ctrl mice. In *Cbx3* KO mice canonical nucleoli (n) were detected, scale bar = 5μm **g-right**: Magnification showing the area of interest with a canonical nucleolus in the *Cbx3* KO mouse (g: granular component; fc: fibrillar centre; dfc; dense fibrillar component). scale bar = 2μm (**h**) rRNA 45S and 18S expression levels in both crypt and villi in the *Cbx3* KO mice (n=3-4mice/group, Student’s *t* test) **(i**) Representative immunofluorescence with anti-lysozyme antibody (red) and Dapi (blue) at the ileal crypt (Scale bar: 20μm) (**j**) Quantification of the lysozyme expression area. (n=6 mice/group, with a total of n=40 field /conditions, Student’s *t* test) (**k**) mRNA expression of sucrase isomaltase in the villi epithelium (n=4-8 mice, Student’s *t* test)

**Extended data Figure 4.**
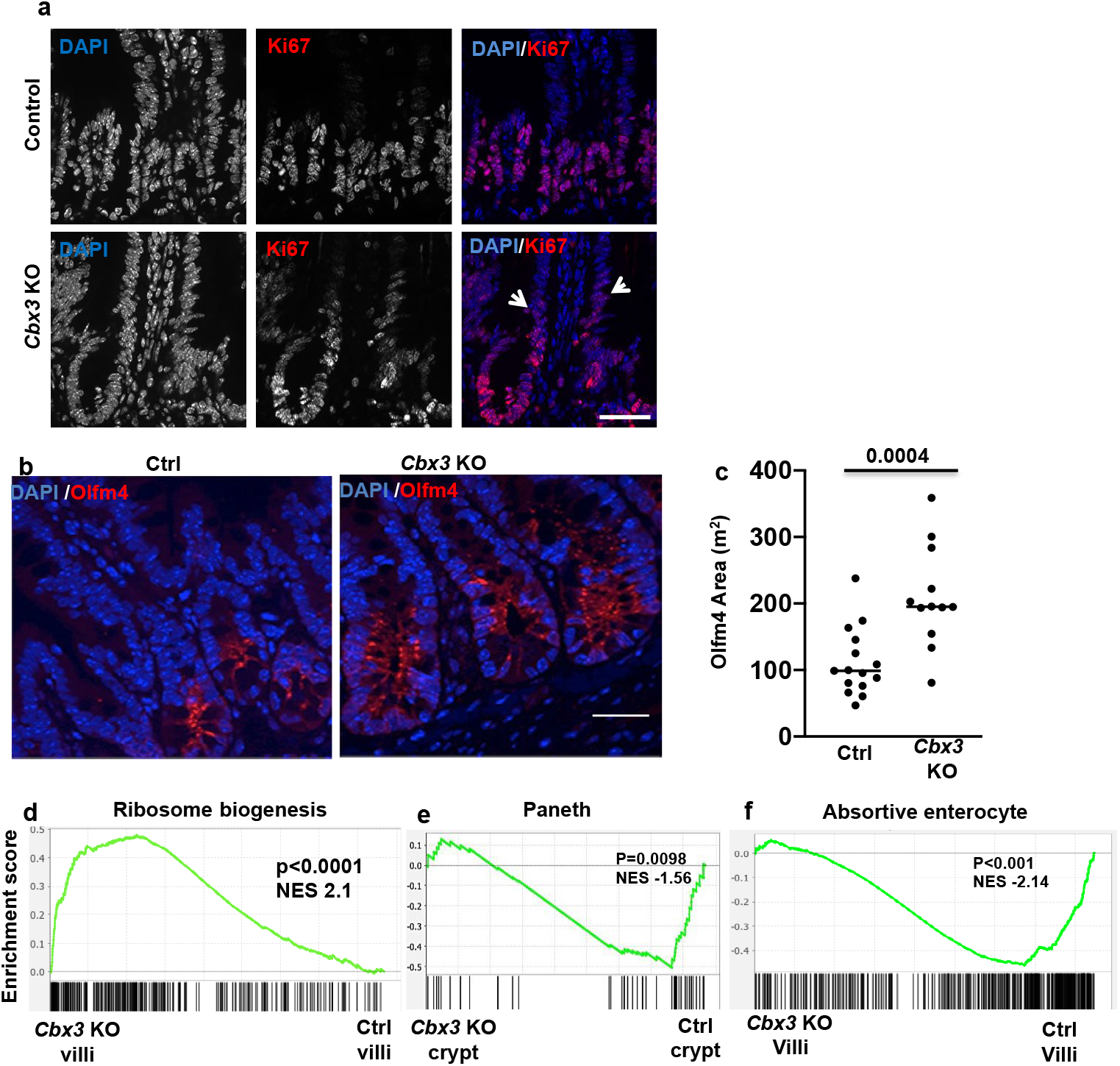
**In (a) Extended KI67 signal at the crypt villi axis in the Cbx3 KO small intestine:** Representative immunostaining of KI67 (red) and Dapi (blue) in control and *Cbx3* KO mice. Arrow head showed aberrant KI67 signal detected at the base of the villi epithelium. Scale bar: 50μm. **In (b-c) Area of detection of the stem cell mark Olfm4** (**b**) Immunofluorescence with olfm4 antibody (red) and Dapi (blue) of ileal crypt sections from Ctrl and *Cbx3* KO mice, (Scale bar: 50μm) (**c**) Quantification of the Olfm4 expression area. Statistical analysis carried out in n=6 mice, measuring a minimum of 20 fields/mice, Student’s *t* test **In (d-f) GSEA analyses:** Significant enrichment of the villi *Cbx3* KO transcriptome with ribosomal biogenesis signature (GOBP_RIBOSOME_BIOGENESIS) In **(e-f)** Inverse correlation between *Cbx3* KO and ctrl (control) mice for Paneth and absortive enterocyte signatures (Haber et al, 2017) at the crypt and villi compartments, respectively. Two-sided nominal *P* values were calculated by GSEA.

**Extended data Figure 5.**
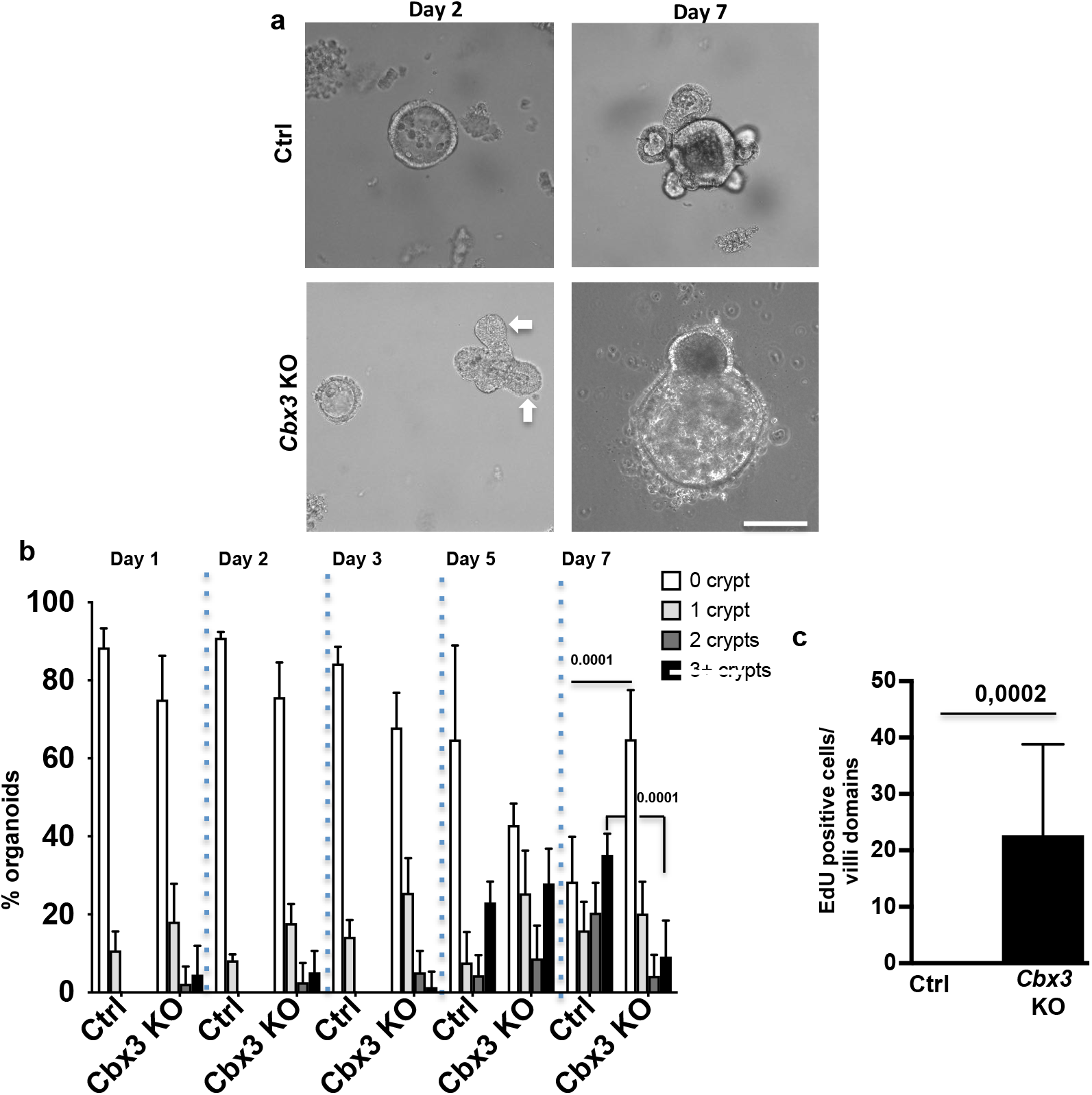
*Ex vivo* enteroid 3D matrigel cultures upon *Cbx3* inactivation: in (a-b) Time course of organoid budding, (**a**) Representative image, taken by transillumination microscopy. Initial increase in budding formation (white arrows) was not maintained overtime. Scale bar: 80μm. (**b**) Evolution of the number of crypts per organoid over time. At day 7, organoid complexicity (i.e. budding crypt potential) was significantly higher in ctrl, as compared to *Cbx3* KO organoids. Counting was carried out on a minimum of 100 organoids, in three different animals, per condition. Statistical analysis carried out by one-way ANOVA. In **(c)** Quantification of the EdU signal in organoids derived from 3 different animals /group, from a total of n=10 organoids/group, Student’s *t* test

Along with this altered proliferative homeostasis, villi from mutant mice were subject to maturation defects, as illustrated by a GSEA analysis, showing a strong association between genes deregulated by *Cbx3* inactivation and genes associated with an intestinal stem cell signature, while RT-qPCR reactions revealed increased expression of the stemness markers Ascl2 and Olfm4 in the mutant mice (**Figures 2 d-e).** Along the villus epithelium of the *Cbx3* KO mice, we also noted expression of nucleolin, a marker of the nucleolus, and electron microcopy studies provided evidence for the presence of nucleoli at the upper part of the villi, with detectable granular components and fibrillar centers **(Figures 2f-g)**. This was in sharp contrast with the expected decline of nucleolin expression along the crypt-villus axis (**Figures 2f-g**), a consequence of the progressive dilution of ribosomes in the post-mitotic cell populations normally thriving on ribosomes inherited from the progenitor cells^18^. Ectopic production of nucleolar organelles was finally documented by an increased 18s rRNA production observed at both crypts and villi, in association with an enrichment in genes involved in ribosomal biogenesis **(Figures 2h and Extended data Figure 4d).** Thus, the homeostatic repression of nucleolar organelle occurring during epithelial cell maturation was lost. Finally, the production of mature lineages in the *Cbx3* KO mice was affected on both absorptive and secretory lineages. Likewise, GSEA analysis of the RNA-seq data from both compartments showed alterations in the Paneth and enterocyte genes expression programs (**Extended data Figures 4e-f**), and we noted a marked defect in the expression of lysozyme, a Paneth cell marker, and of Sucrase Isomaltase, an absorptive enterocyte differentiation marker (**Figures 2i-k**). Overall, these data are indicative of an extensive deregulation in the control of cell proliferation and in the production of mature lineages upon loss *Cbx3* activity.

### HP1γ limits usage of poor splice sites

We next sought to define functions for HP1γ on RNA homeostasis in the gut epithelium. Mass spectrometry revealed that HP1γ interactants were highly enriched in component of the spliceosome (**Extended data Figure 6a-c and Supplementary Data Table 5**). Among them, we identified members of the catalytic step 2 spliceosome (also called U2-type spliceosomal complex C) and splicing factors essential in splice-site recognition, such as the Ser/Arg-rich (SR) proteins family (**Extended data Figure 6c**).

**Extended data Figure 6.**
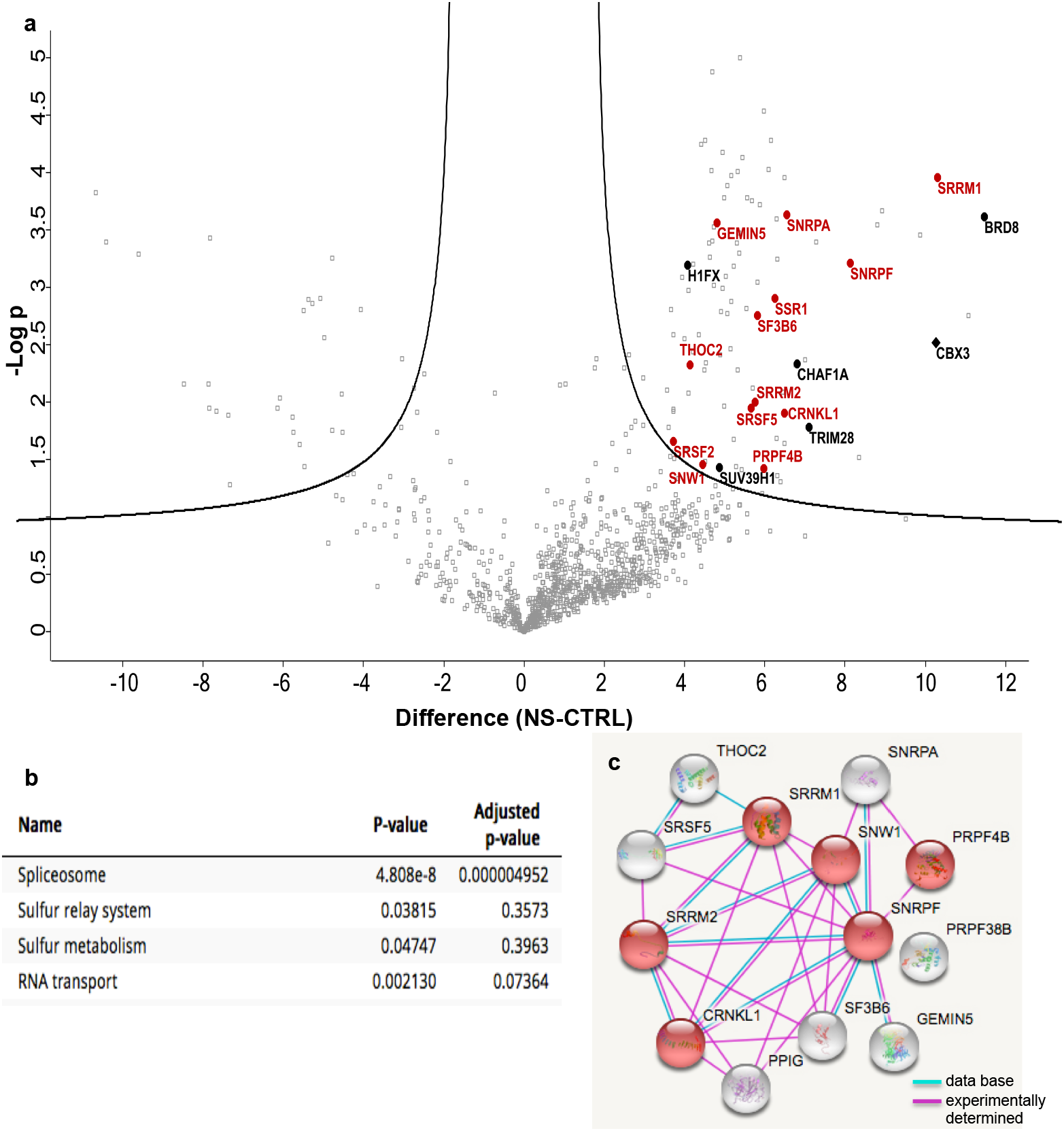
Proteomic analysis: **(a)** Volcano plot representation of proteins immunoprecipitated with anti-HP1γ or IgG (CTRL) antibodies in HeLa cells and identified by mass spectrometry in 3 independent experiments. X-axis reports the difference of the average of the logarithm of Label Free Quantification (LQF) intensities between IgG control and with anti-HP1γ immunoprecipitates. Y-axis reports the negative logarithm of t-test p value. In black: classical HP1 interactants In red: spliceosome interactants. (**b**) KEGG-pathway analysis on the molecular interactants identified by mass spectrometry (a total of 96 hits). (**c**) Predicted interactions in mass-spectrometry hits matching the KEGG-pathway “spliceosome”. Modules in red map to Catalytic step 2 spliceosome (GO:0071013, FDR 4.61e-08).

This was consistent with earlier reports on the role of HP1 γ in the regulation of pre-mRNA splicing^5,19^. Likewise, analysis of the RNA-seq data with the Multivariate Analysis of Transcript Splicing (rMATS) pipeline confirmed that splicing was extensively impacted upon HP1 γ inactivation. Significant variations in splicing included a trend towards increase intron retention in the small intestine but not in the colon, with increased and decreased inclusion of alternative exons were observed (**Figure 3a and Supplementary data Tables 6-8,** significant (FDR<0.05) alternative splicing events in crypt, villi and colon epithelia). The absence of a defined pattern in the impact of HP1γ on splicing prompted us to examine whether these numerous splicing events would result from noisy splicing^20^, as related in cancer and neurodegeneration ^21–24^

**Figure 3.**
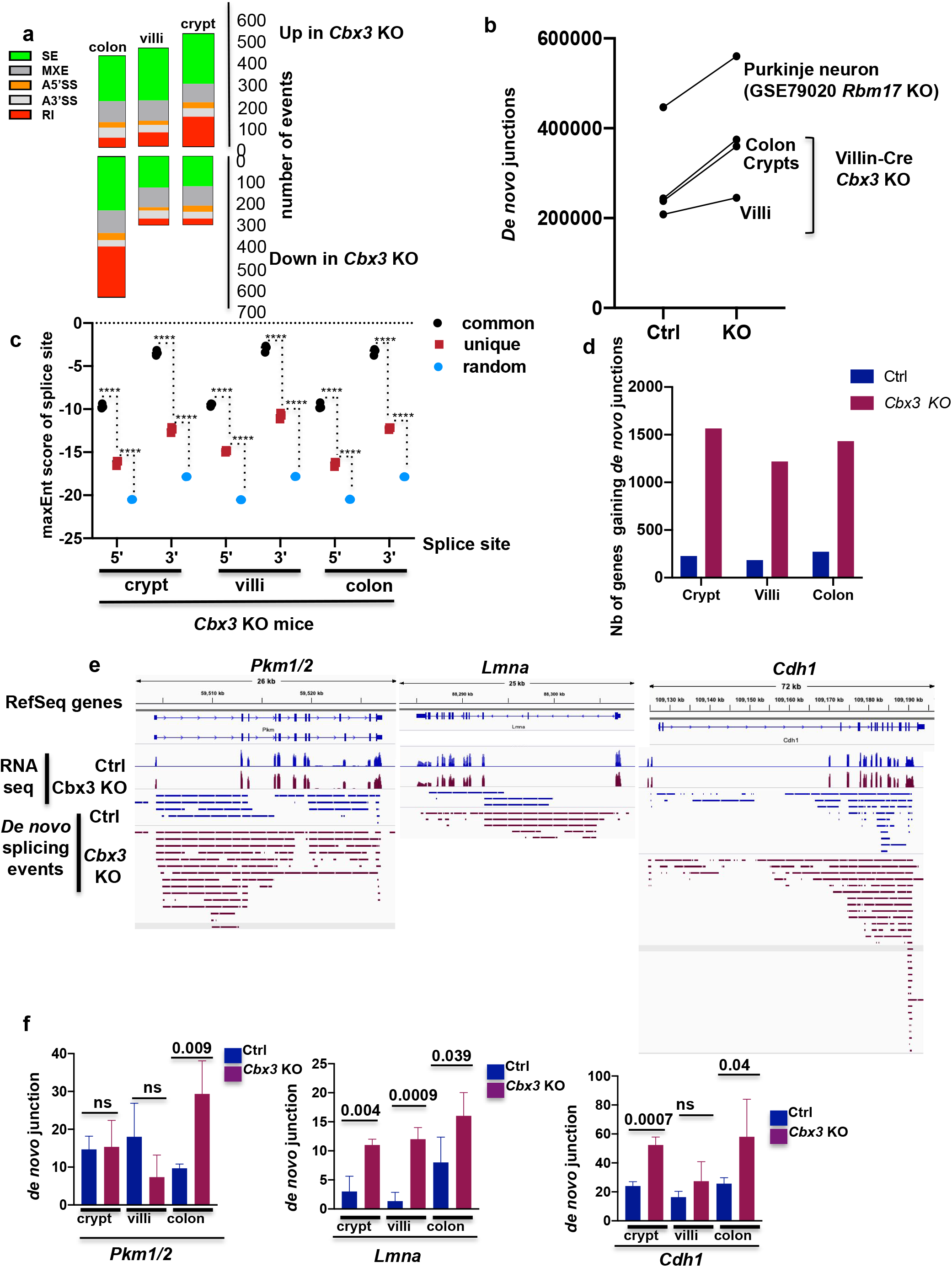
Impact of HP1γ deficiency on RNA splicing *in vivo*. **(a)** Types and quantity of splicing alterations detected in the colon, villus, and crypt epithelia. Types of splicing alterations are color coded as indicated. **SE**: skipped exons, **MXE**: mutually exclusive exons, **A3’SS** and **A5’SS**: alternative 3’/5’ splice sites, **RI**: retained introns. (**b**) Quantification of *de novo* junctions in control (Ctrl) and *Cbx3* KO (KO) conditions in crypt, villi and colon epithelia after consolidating the 3 RNA-seq replicates from either ctrl or *Cbx3* KO mice in each tissue. *De novo* junctions were defined as junctions not annotated in the mm9 version of the mouse genome and not present in both WT and mutant samples. P-Value was calculated with a paired t-test, n=3. Quantification of *de novo* junctions in *Rbm17* KO mice in Purkinje neurons (GSE79020) is shown (**c**) Consolidated maxEnt score of the *de novo* sites identified with the approach described in (**b**) (red), compared to the score obtained with annotated junctions and that obtained with randomly selected sequences (black and blue, respectively). Values shown are from *Cbx3* KO crypt, villi, and colon (n=3 mice, Ordinary one-way Anova ****p<0.0001). (**d**) *De novo* junctions were quantified at genes that, transcriptionally, were affected less than 2-fold by *Cbx3* KO inactivation. For each indicated condition, we counted genes gaining *de novo* junction 2-fold or more (p Val<0.05). (**e**) *De novo* junctions in colon (PKM and CDH1) or villi (LMNA) were visualized with a genome browser in the neighborhood of the indicated genes. Bars represent the chromosome region located between the 3’ and the 5’ splice sites for each *de novo* junction. (**f**) Bar graphs show the number of *de novo* junction detected at the corresponding genes at crypt, villi and colon ((n=3 mice for each condition, Student’s *t* test).

To that end, we identified unannotated splice junctions present in only one of the two experimental conditions (either WT and Cbx3 KO) for each tissue. Abundance of these junctions that we henceforth will refer to as *“de novo* was significantly increased upon *Cbx3* inactivation (**Figure 3b**). This increase was in a range similar to that observed in cerebellar Purkinje neurons depleted of *Rbm17,* an RNA binding protein reported to repress cryptic splicing usage^23^ (**Figure 3b**).

Evaluating the quality of the consensus splicing donors and acceptors with a dedicated software revealed that, in average, the *de novo* junctions scored lower that annotated junctions, but higher than random sequences (**Figure 3c)**. This suggested that inactivation of HP1 γ promoted usage of poor consensus splice sites, and possibly increased the opportunity range of splicing.

Finally, examining splicing noise induced by *Cbx3* inactivation on a gene-per-gene basis further documented that, at a large majority of genes not affected at their expression level, the number of active splice sites was significantly increased (**Figure 3d and Supplementary data Table 9 for the list of genes)**. These genes included regulators of gut homeostasis. In particular, we noted that the *Pkm* gene, producing both Pkm1 and Pkm2 mRNAs by alternative splicing, the latter safeguarding against colitis^25^, showed an extensive increase in *de novo* splicing in the colon (**Figures 3e and f, left panels**). Likewise, the *Cdh1* gene encoding E-cadherin product, essential for the epithelial barrier function, and subject to premature termination by alternative splicing^26^ was affected by *Cbx3* inactivation at both crypts and colon (**Figures 3e and f, middle panels**). Finally, we observed a particularly strong impact at the *Lmna* gene with an average 9-fold increase in *de novo* junctions at the villi (**Figures 3e and f, right panel**).

### Control of the progerin splice variant by HP1γ

The biological significance of splicing noise has been poorly characterized in mammals, prompting us to investigate whether the increased usage of *de novo* splice-junctions at the *Lmna* gene in villi lacking HP1γ could result in the production of progerin. Progerin is a splice variant of the Lmna pre-mRNA responsible for the Hutchinson Gilford Progeria Syndrome (HGPS)^27–28^. In this syndrome of premature ageing, production of this splice variant is facilitated by a genomic mutation increasing usage of a progerin splice site that in normal cells is used at low yield upon usage of a poor-consensus splice site^29^.

We thus applied a taqman assay to the detection of laminA and progerin splice isoforms in the mouse gut epithelium **(Figures 4a-b)**. *Cbx3* inactivation resulted in a significantly increased occurrence of the progerin-specific splicing event in the villus epithelium **(Figure 4b)**. Comparison with colon intestinal epithelium from the *Lmna* G609G HGPS mice revealed that progerin-specific splicing events induced by *Cbx3* inactivation remained less frequent than in cells carrying the *Lmna* G609G mutation, in coherence with the increased strength of the progerin 5’SS provided by the mutation^29^(**Figures 4b**). Moreover, sequencing of the PCR end-products confirmed that the splice events detected in *Cbx3* KO mice were identical to the splicing event generated by the HGPS mutation (**Figure 4c**). Levels of the canonical Lamin A splicing events were also increased in both crypts and villi in the mutant mice (**Figure 4a**), while expression of the *Lmna* gene remained unaltered as documented by the transcriptome analysis (**Figure 3e, middle panel**). Finally, the detection of progerin transcript in the small intestine of the *Cbx3* KO mice was correlated with the production of progerin protein at both villi and crypt epithelia, as visualized by western (**Figures 4d**). In these assays, we used a well-characterized anti-progerin monoclonal antibody, readily detecting the progerin protein in the colon epithelium of HGPS mice (**Extended data Figure 7)**. Of note, the absence of progerin signal in *Cbx3* fl/fl not harboring an inducible Cre-recombinase allowed to rule out an effect of the tamoxifen induction on progerin production (**Figure 4d**). Accumulation of progerin protein in crypts and villi was further documented by immunocytochemistry (**Extended data Figure 8).** This approach revealed that progerin was principally detected at the immediate progeny of ISC (above +4 position, **Figures 4e-f**), thus evidencing for a gradient of progerin expression along the crypt-villi axis in *Cbx3* KO mice. *In vitro* crispr/Cas9-mediated *Cbx3* knockout (KO) in enterocytic TC7 cells confirmed the production of progerin transcripts, and accumulation of nucleocytoplasmic progerin protein (**Extended data Figures 9a-b**).

**Figure 4:**
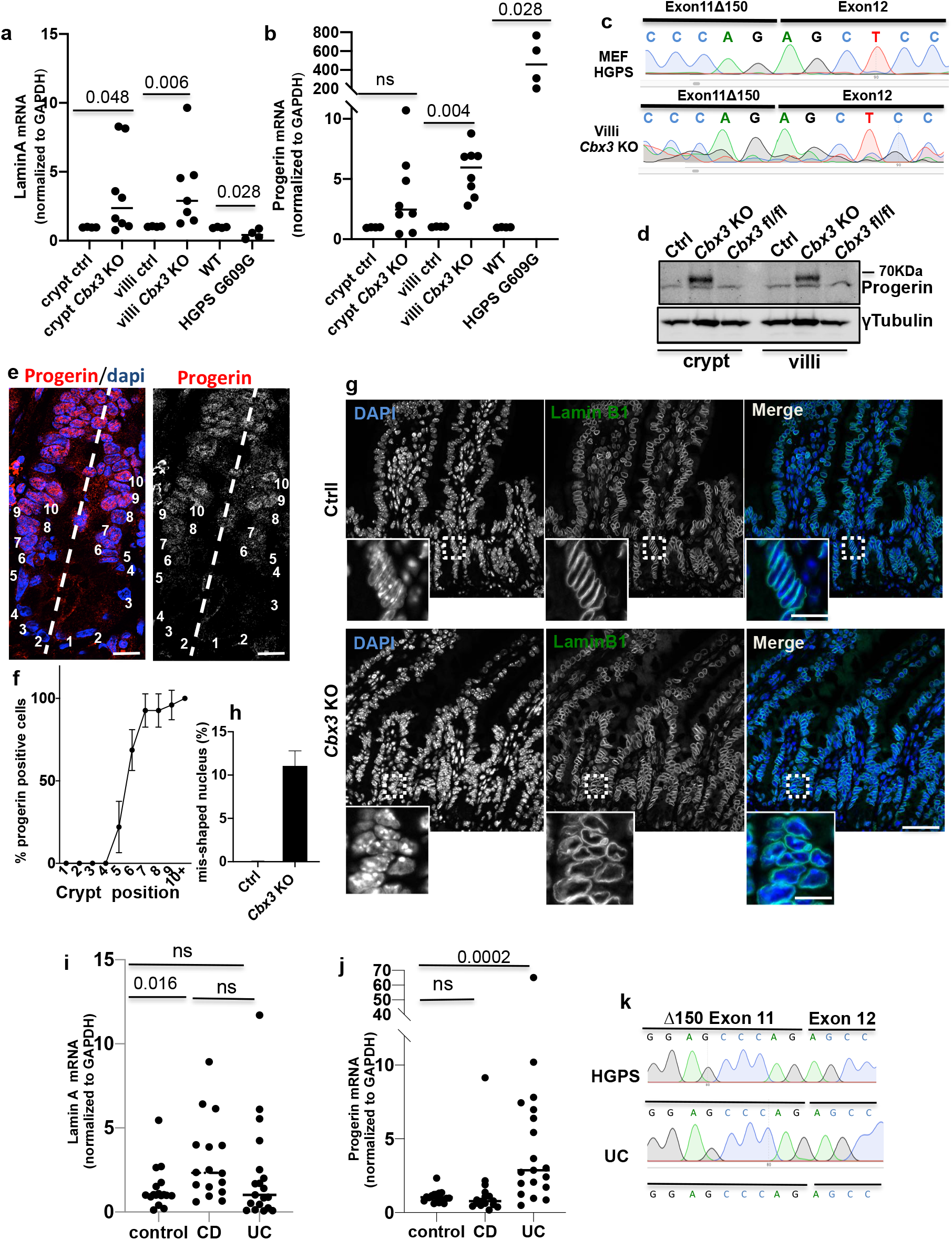
Analysis of progerin production in the gut epithelium. (**a,b**) Taqman Assays laminA and progerin in ctrl (n=4), *Cbx3* KO (n=8), WT and HGPS G609G (n=4mice/group), Mann-Witney U test (**c**) examples of sequencing data of the Taqman PCR end products in villi *Cbx3* KO samples, mouse embryonic fibroblast (MEF) derived from G609G mice used as control (**d**) Immunoblot with anti-progerin monoclonal antibody in epithelium lysates derived from Ctrl, *Cbx3* KO mice and as internal control, *Cbx3 fl/fl* not expressing the Cre recombinase treated with tamoxifen, (**e-f**) Immunofluorescence with anti-progerin antibody (red) and Dapi (blue) and percentage (%) of progerin expressing cells according to the position along the ileal crypt axis (*Cbx3* KO n=23 sections, n=3 mice). Values are represented by the mean with SD. Scale bar: 20μm (**g**) Immunofluorescence with anti-Lamin B1 antibody (green) marking the nuclear envelope (Scale bar: 50μm, Insert: 15μm (**h**) Percentage of cells with misshaping nucleus. n=5 mice in each group, counting in ctrl=4757 nucleus and *Cbx3* KO=29753 nucleus counted, respectively). Student’s *t* test (**i,j**) Taqman Assays laminA and progerin in control (n=17), Crohn disease (CD, n=16) and Ulcerative colitis (UC,n=19) populations, cDNA were extracted from colon biopsies. Mann-Witney U test (**k**) example of sequencing data from the Taqman PCR end products in one UC patient, Human fibroblast from HGPS patient was used as positive control, UC= Ulcerative colitis.

**Extended data Figure 7.**
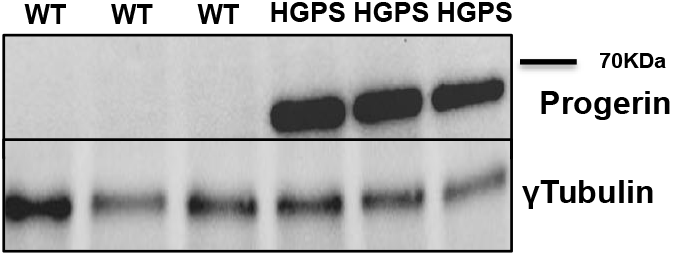
Progerin expression detected by Immunoblot in colon epithelium lysates derived from 12 months old WT and heterozygote HGPS G609G mice (n=3 mice in each group)

**Extended data Figure 8.**
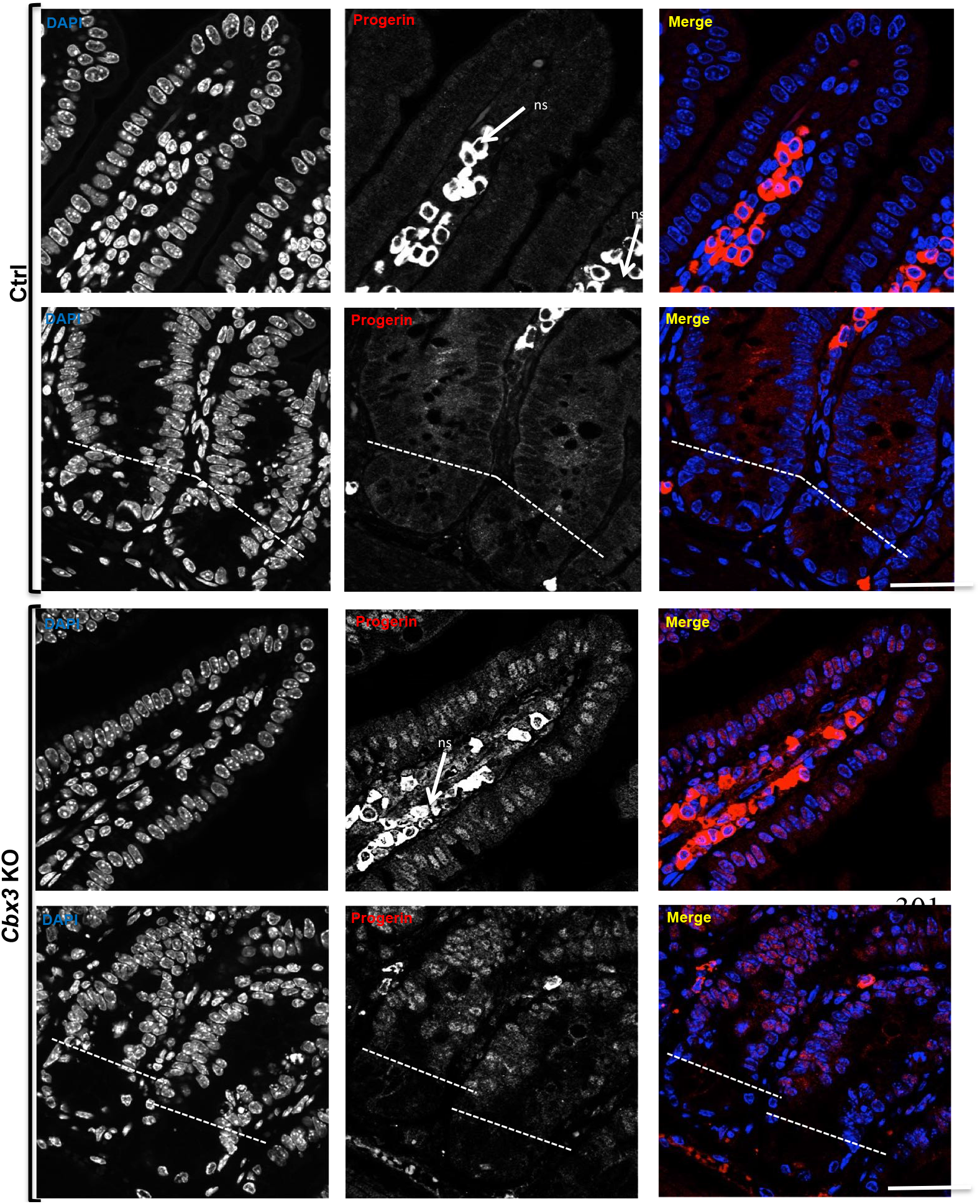
Representative immunostaining of progerin in the ileon crypt and villi epithelia: progerin signal (red) is detected in the nucleus (Dapi, blue) *ns= non specific labeling (detectable with secondary antibody alone), dashed line showing the stem cell compartment at the intestinal crypt (position O to +4) (Scale bar: 80μm)

**Extended data Figure 9.**
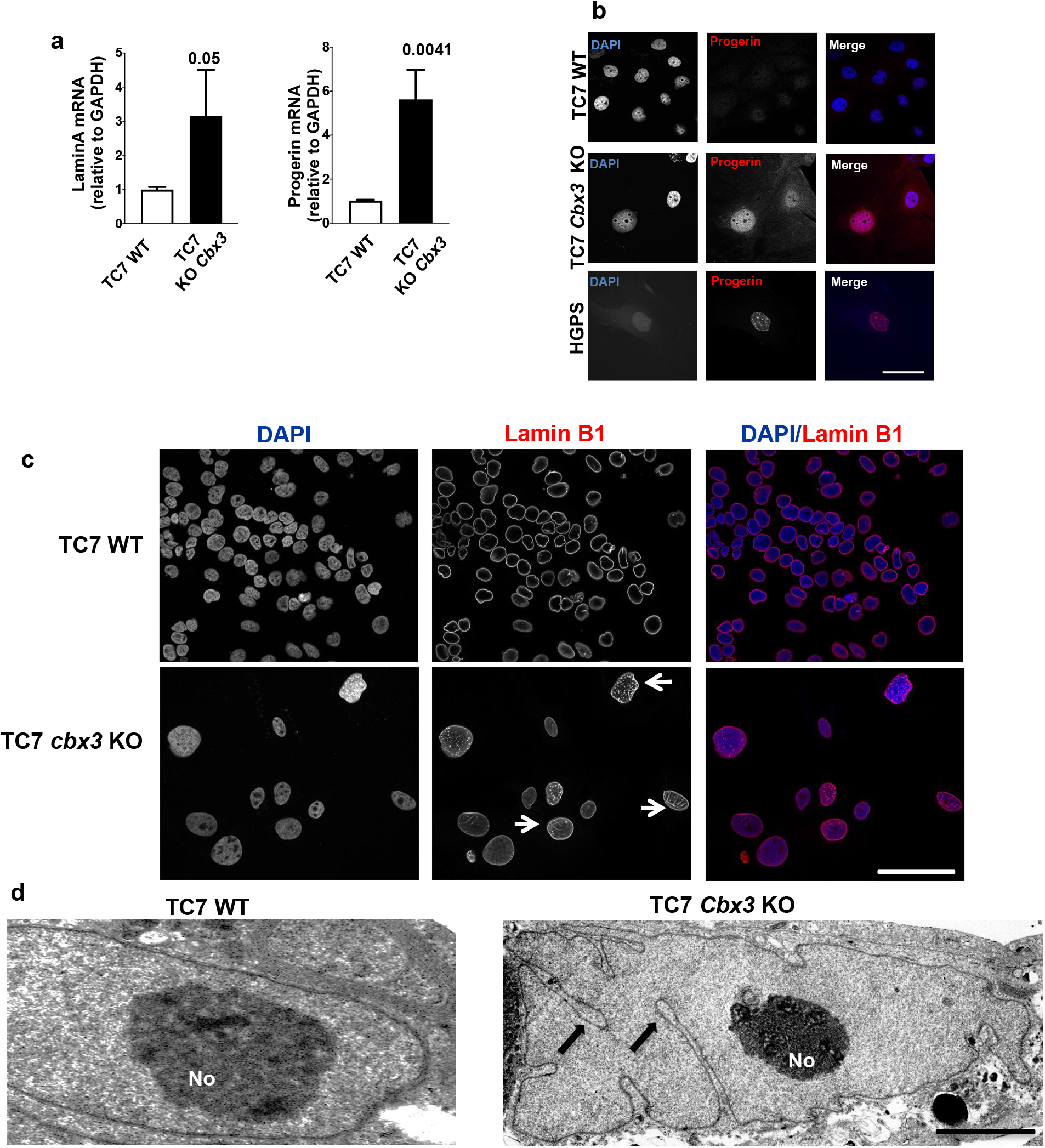
*Cbx3* inactivation leads to a laminopathy with progerin production in the Crispr/Cas9 *Cbx3* enterocytic cell line TC7 (TC7 *Cbx3* KO). (**a**) RT-qPCR lamin A and progerin in the WT and Crispr/Cas9 *Cbx3* enterocytic cell line TC7 (TC7 *Cbx3* KO). Values are represented by the mean with SD of 3 independent experiments, Student’s *t* test (**b**) Representative immunofluorescence with anti-progerin antibody (red) and Dapi (blue) in WT, TC7 *Cbx3* KO and human HGPS fibroblast **(c)** Immunostaining with anti-lamin B1 antibody: laminB1 signal (red) reveals mis-shaping at the nuclear envelop (white arrow) reminiscent of progeria cells (Scale bar: 50μm).(**d**) Transmission Electron Microscopy (TEM) on TC7 cells (WT and *Cbx3* KO). The nuclear envelop shows multiple invaginations in TC7 *Cbx3* KO cells (black arrows). No = Nucleolus. (Scale bar: 2μm WT, 5μm *Cbx3* KO)

Finally, we examined whether levels of progerin produced upon *Cbx3* inactivation were sufficient to induce toxicity to the nuclear lamina. Observation of the small intestine of *Cbx3* KO mice revealed a misshaping of the nuclear envelope in a fraction of the epithelial cell upon laminB1 immunostaining (**Figures 4g-h**). Similar defects were also observed *in vitro* upon crispr/Cas9-mediated *Cbx3* knockout (KO) in the TC7 enterocytic cell line, resulting in deep invaginations of the nuclear membrane detectable by electron microcopy, reflecting a laminopathy phenotype (**Extended data Figures 9c-d)**. Overall, these data showed that HP1γ defect increased the opportunity range of laminA mRNA splice variants, thus leading to the production of progerin in the absence of HGPS mutation in the gut epithelium.

Finally, we investigated whether progerin was also produced in colonic tissue from UC patients, that, as shown **Figures 1a-b,** expressed reduced levels of HP1γ. cDNA were extracted from colon biopsies in non-inflamed zones of healthy individuals (n=17), UC patients (n=19), or Crohn’s disease (CD) patients (n=16) with colonic involvement **(Supplementary Data Table 10 for detailed description of the population)**. Lamin A and progerin mRNA expressions were determined by taqman assays and accurate transcript detections by sequencing of the PCR end-products. In UC patients, levels of progerin-specific splicing events by taqman assays were significantly up-regulated (p Val= 0.0002), (**Figures 4j-k**), while production of the laminA splicing product was not (**Figure 4i**). No correlation was detected with between progerin mRNA expression and patient age (**Extended data Figure 10a).** Inversely, in CD patients, progerin-specific splicing was unaffected (**Figure 4j**), while the lamin A-specific splicing product was up-regulated (p Val=0.016) (**Figure 4i**), this increase mainly concerning the younger CD patients less than 45 years old, as shown by linear regression analysis **(Extended data Figures 10b-c)**. Thus, progerin transcript emerges as a potentiel marker of UC, while our observations also emphasize the predictive value of the *Cbx3* KO mouse model in this disease.

**Extended data Figure 10.**
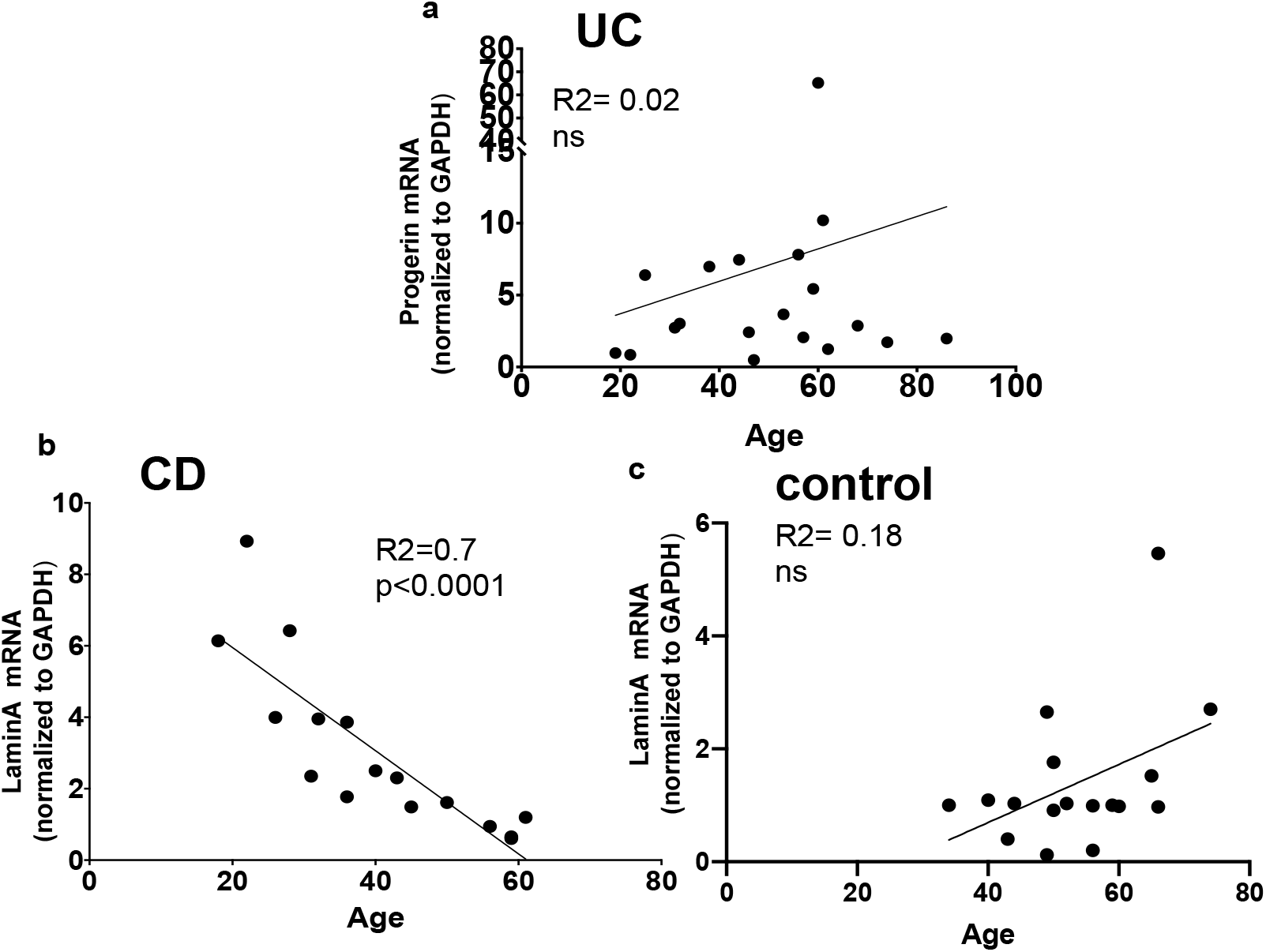
In **(a**) Simple regression analyses for progerin mRNA expression in UC (ulcerative colitis) patients relative to the age In **(b-c**) Simple regression analyses of laminA mRNA expression relative to the age in (**a**) CD (Crohn’s disease) and (**c**) control patients. The pvalue corresponds to the linear regression of laminA expression vs age

## Discussion

This work seminally defined HP1γ as a mechanism safeguarding RNA splicing accuracy in the gut epithelium, limiting the impact of naturally occurring non-canonical splicing events. Non-canonical splicing activity is frequently referred to as “splicing noise”^20^. While noisy splicing in non-mammalian animal model may promote diversity in mRNA splice variants, with possible physiological functions^30–31^, in mammals, its significance remains elusive, and seen as a pervasive erroneous splicing activity, possibly involved in genome evolution^32^. As a matter of fact, this phenomenon has been linked to human pathology, with global activation of noncanonical splicing sites, as detected in cancer and neurogenerative diseases ^22–24,33^. Only few RNA binding proteins, including Rbm17, hnRNP and TP43^23–24^, were shown to dampen cryptic splicing genome-wide and our study highlights a fundamental role for HP1 in this process in the gut epithelium. In this context, HP1γ deficiency was sufficient to induce the production of progerin, a normally repressed splice variant. To our knowledge, HP1γ is the first identified *in vivo* mechanism regulating progerin production in the absence of the HGPS mutation. The effects of HP1γ inactivation on splicing was accounted for by an apparently general reduction in the precision of splicing. Specifically, junction reads were increasingly detected at non-annotated splice junctions, and at sites with very poor donor or acceptor consensus sequences. While our understanding of this decreased stringency in splice site selection is still incomplete, we note that our proteomic approach to HP1γ molecular partners detected numerous splicing factors ensuring the usage of only genuine splice sites. These included components of the major (U2-dependant) spliceosome as well as auxiliary splicing factors essential for spliceosome recognition of the consensus sequence. Among these, the Ser/Arg-rich (SR) proteins, were previously shown to modulate usage of the splice sites implicated in the production of progerin in the absence of the HGPS mutation, possibly by interfering with exonic splicing enhancers (ESE) involved in proper exon recognition by SR proteins^34,29^. We thus propose a model in which the chromatin-associated HP1 γ participate in the precise co-transcriptional recruitment of regulators of pre-mRNA splicing. The reliance of HP1 γ on auxiliary regulators of splicing may also be illustrated by its regional impact on splicing, possibly as a function of the splicing regulators available in each tissue or cell type. In particular, we noted a gradient of progerin expression along the crypt-villi axis, leaving out the ISC compartment. The latter suggests a modulation of splicing possibly linked to the cellular developmental state, a notion further supported by the HGPS-iPSCs model, in which progerin was only observed upon cell commitment^35^. While our data mainly points toward an impact of HP1 γ via spliceosome recruitment, an involvement for HP1γ in the degradation of defective splicing products may also be considered, as suggested by the implication of the yeast HP1 homolog in escorting pericentromeric transcripts to the RNA decay machinery^36^.

Before a role for HP1 was described in splicing, this protein was mainly known as a mediator of heterochromatin-dependent silencing, repressing transcription at repeated DNA sequences, including rDNA loci, and at a subset of gene promoters ^8,16,37–40^. This repressive function of HP1γ was found to also have a clear impact on the biology of the small intestine, participating in the regulation of the nucleolar organelles, and in the silencing of genes controlling proliferative homeostasis. These observations illustrate the homeostatic functions of HP1γ involved in essential activities in both heterochromatin and euchromatin. The previously described role of HP1 in tissue longevity across the species^40–41^ may be largely rooted in this polyvalence.

While no human disease has been clearly linked to HP1 gene mutations, decreased HP1 expression was reported in syndromes of accelerated aging, including the Werner syndrome and HGPS^42–44^. We similarly identified a potent reduction of HP1 expression in UC patients as well as in mouse model relevant for this disease. From our *Cbx3* KO mouse model, we concluded that HP1γ deficiency may support aberrant pathways nurturing both inflammation and splicing defects at key homeostatic genes. Likewise, the increased detection of progerin in UC colon tissue prompts us to speculate that, alike what we observed in the *Cbx3* KO mice, the lamin A splice variant is only the “tip of the iceberg”, indicative of a more extensive disturbance in RNA splicing precision endured by UC patients. Still in its infancy with regard to intestine disease^45–47^, the identification of mechanisms relying on RNA splicing dysfunctions should transform our understanding of the functional decline in the gut epithelium of IBD patients.

## Methods

### Mouse models

*Cbx3*^*fl*/fl^ mice (provided by Dr Florence Cammas, IRCM, Montpellier, France) were crossed with Villin-CreERT2 mice (provided by Dr Cohen-Tannoudji, Institut Pasteur, Paris, France) to produce the Villin-creERT2:*Cbx3-/-* mouse model (this study). Heterozygote *Lmna^G609G/G609G^* mice (here referred as HGPS mice) were provided by Dr Maria Eriksson (Karolinska Institutet, Sweden). Mice were fed by a standard diet (SD) rodent chow (2018 Teklad Global 18% Protein Rodent Diet, Harlan) composed of 60% carbohydrate fed *ad libitum*. Tamoxifen (0.5mg/g) diluted in 20% clinOleic acid was administrated by oral gavage, at 3 doses every 5 days as described^48^. Control mice received 20% clinOleic acid alone by oral gavage. Additional controls using *Cbx3*^*fl*/fl^ mice that do not express the Cre recombinase were identically treated with tamoxifen. All the experiments using Villin-creERT2: *Cbx3-/-* mouse model were performed with mice 2-3 months of aged. BrdU (Sigma) was injected intraperitoneally (i.p.) at 100 μ g/g animal body weight, 1h prior to sacrifice.

### Patients and Biopsy Specimens

All patients were followed in the Department of Gastroenterology (hôpital Beaujon, Paris). The protocol was approved by the local Ethics Committee (CPP-Ile de France IV No. 2009/17, and No2014-A01545-42) and written informed consent was obtained from all patients before enrollment. Colonic pinch biopsy allowing the extraction of the epithelial layer was obtained during endoscopic investigation in non-inflamed areas to allow a comparison with healthy tissues in the control population. Biopsies were performed in the right or the left colon or in case of cancer at least 10 cm away from the cancer site. For the immuno-histological study, 26 patients were included, with 10 healthy control and 16 UC patients (detailed of the population provided in **Supplementary data Table 1**). For the transcriptional study, 56 patients were included, with 17 healthy controls, 19 UC and 16 CD patients with colonic involvement (detailed of the population provided in **Supplementary data table 10**). Total RNA was extracted from human biopsies with RNAble (Eurobio) and quantified using a ND-1000 NanoDrop spectrophotometer (NanoDrop Technologies). Reverse transcription of total mRNA was performed using M-MLV RT (Invitrogen) according to the manufacturer’s recommendations.

### Cell cultures

Human dermal fibroblasts patients who carried the HGPS p.Gly608Gly mutation were obtained from the Coriell Cell Repository. Caco2 (TC7 cells) were used to generate the CRISPR/Cas9-mediated *Cbx3* cell line. The pSpCas9(BB)-2A-GFP (PX458) vector expressing Cas9 endonuclease (gift from Feng Zhang, Addgene plasmid # 48138) was linked with a single-guide RNA (sgRNA) designed specifically for *Cbx3* gene. Two Sequence guides (GAAGAAAATTTAGATTGTCC and GAATATTTCCTGAAGTGGAA) were defined by ZiFiT Targeter Version 4.2 software. Insertion of the sequence guide was performed in the BbsI restriction site of the PX458 vector and check by sequencing. Transfection in TC7 cells was performed by lipofectamine 2000 and singles clones for each sequence guide were selected by FAX according to the GFP signal.

### Immunofluorescence

Intestine (ileum or colon) was collected and washed with PBS at 4°C and cut in pieces about 5 mm. Intestinal fragments were fixed with formalin overnight at 4°C. Once fixed, intestinal fragments were included in paraffin blocks. Paraffin sections were done in a microtome Leica RM2125 RTS, with a thickness of 4μm. Subsequently, the deparaffinization and rehydration of the samples was carried out by immersion in Xylene (2×10 min), absolute ethanol 5 min, 90% ethanol 5 min, 70% ethanol 5 min and distilled water (2×5 min), all at R.T. Finally, the antigen was unmasked using the EDTA boiling technique for 30 min at 95°C, followed by 20 min at R.T. All samples were sequentially treated with 0.1 M glycine in PBS for 15 min, 3% BSA in PBS for 30 min and 0.5% Triton X-100 in PBS for 2h (mouse tissue). In case of nucleolin staining, tissue sections were incubated during one minute at RT with proteinase K 0.05mg/ml, followed by a wash with glycine 2mg/ml during 15 minutes at RT. They were then incubated with primary antibodies overnight at 4 °C, washed with 0.05% Tween-20 in PBS, incubated for 1h in the specific secondary antibody conjugated with Alexa 488 or Cy3 (Jackson, USA), 15 min with DAPI (1μg/ml), washed in PBS and mounted with the antifading medium VECTASHIELD® (Vector laboratories). Anti-HP1γ (2MOD-IG6, Thermo Scientific), HP1β (1MOD-1A9, Thermo Scientific), HP1α (2H4E9, Novus Biologicals), KI67 (ab16667, Abcam), Olmf4 (D6Y5A, Cell Signalling), Brdu (MA3-071, Thermo Scientific), laminB1 (ab65986, Abcam), Progerin (13A4DA, sc-81611, Santa Cruz), γTubulin (4D11, Thermo Scientific), and Nucleolin (ab22758, Abcam) were used as primary antibodies. Nuclei were stained using 4,6-diamidino-2-phenylindole (DAPI, 62248, Thermo Scientific).

Microscopy images were obtained with a ZEISS Apotome.2 (Zeiss, Germany), structured illumination microscope, using a 63× oil (1.4 NA) objective. To avoid overlapping signals, images were obtained by sequential excitation at 488 and 543 nm in order to detect A488 and Cy3, respectively. Images were processed using ZEISS ZEN lite software. The quantitative analysis of the immunofluorescence signal was performed on ImageJ. The values are represented with Mean fluorescence intensity, relatives to control samples.

### Tissue processing for intestinal epithelial cells isolation and organoids culture

The technique was adapted from *Nigro* et al, 2019^49^. Small intestine or colon were collected and washed with PBS at 4°C and cut in pieces of about 5 mm in length. For epithelial isolation, intestinal fragments were incubated 30 min at 4°C in 10mM EDTA after which they were transferred to BSA 0,1% in PBS and vortexed 30-60 s. The supernatants (containing the epithelial cells) were filtered with a 70μm cell strainer. At this step, crypts went through the cell strainer and villi were retained on it. Crypt and villus fractions were then centrifuged separately and the pellets were frozen in liquid nitrogen until processed. The quality of the separation was accessed by the expression of selective expression of stemness markers in the crypts but not villi, as confirmed by RNA sequencing. For organoid production, crypt pellet was disaggregated and cultured in Matrigel as described ^49^. EdU staining was performed using Click-iT™ EdU Cell Proliferation Kit for Imaging (Thermo fisher), following manufacturer indications.

### Transmission electron microscopy

In TC7 cells, Transmission Electron Microscopy (TEM) was performed by cryofixation/freeze substitution method. Cells were fixed overnight at 4°C with 3.7% paraformaldehyde, 1% glutaraldehyde, in 0.1 M cacodylate buffer. After fixation, cells were pellet and contrasted with osmium tetroxide (OsO4) and uranyl acetate. Then, cells were included in freeze-substitution medium during at least 3 days at −80°C, dehydrated in increasing concentrations of methanol at −20 °C, embedded in Lowicryl K4 M at −20 °C and polymerized with ultraviolet irradiation. Ultrathin sections were mounted on nickel grids, stained with lead citrate and uranyl acetate and examined with a JEOL 1011 electron microscope.

Intestinal tissue fragments around 500mm were fixed overnight at 4°C with 3.7% paraformaldehyde, 1% glutaraldehyde in 0.1 M cacodylate buffer overnight at 4°. Small tissue fragments were washed in 0.1 M cacodylate buffer, dehydrated in increasing concentrations of methanol at −20 °C, embedded in Lowicryl K4 M at −20 °C and polymerized with ultraviolet irradiation. Ultrathin sections were mounted.

### Real time quantitative PCR analysis

Total RNA was extracted using Trizol (TR-118, Molecular Research Center, Inc.) following the manufacturer’s instructions and DNAse treatment. RNA samples were quantified using a spectrophotometer (Nanodrop Technologies ND-1000). First-strand cDNA was synthesized by RT-PCR using a RevertAIT H Minus First Strand cDNA Synthesis kit (Thermo Scientific). qPCR was performed using the Mx3005P system (Stratagene) with automation attachment. For progerin and laminA detection, real-time PCR amplification was carried out with the TaqMan” Gene Expression Master Mix (life technologies) using predesigned primers for mouse GAPDH (Mm99999915_g1), mouse lamin A that do not recognized progerin or DNA (Assay ID : APGZJEM), mouse progerin (F: ACTGCAGCGGCTCGGGG R: GTTCTGGGAGCTCTGGGCT and probe: CGCTGAGTACAACCT), human lamin A (F: TCTTCTGCCTCCAGTGTCACG R: AGTTCTGGGGGCTCTGGGT and probe ACTCGCAGCTACCG), human progerin (F: ACTGCAGCAGCTCGGGG R: TCTGGGGGCTCTGGGCTCCT and probe CGCTGAGTACAACCT), human GAPDH (Hs00266705_g1). For SYBRGreen (Takara) based qPCR, the following primers have been used: mouse Ascl2 (F: GGT GAC TCC TGG TGG ACC TA ; R: TCC GGA AGA TGG AAG ATG TC) mouse Olfm4 (F:ATC AGC GCT CCT TCT GTG AT R: AGG GTT CTC TCT GGA TGC TG) mouse TNF-a (F : GATCTCAAAGACAACCAACATGTG R : CTCCAGCTGGAAGACTCCTCCCAG) mouse IL1-β (F :TACAGGCTCCGAGATGAACAAC R : TGCCGTCTTTCATTACACAGGA) mouse IL6 (F : ACTTCCATCCAGTTGCCTTCTT R: CAGGTCTGTTGGGAGTGGTATC) mouse Sucrase isomaltase (F: CGTGCAAATGGTGCCGAATA R: TCCTGGCCA TACCTCTCCAA) GAPDH (F: TGACCACAGTCCATGCCATC; R: GACGGACACATTGGGGGTAG) mouse 45S (F: GAACGGTGGTGTGTCGTT; R: GCGTCTCGTCTCGTCTCACT). Mouse 18S: (F: GATGGTAGTCGCCGTGCC; R: CCAAGGAAGGCAGCAGGC).

### RNA sequencing and bioinformatic pipelines

Total RNA was extracted from control and *Cbx3* KO epithelial cells from the small intestine (crypts and villi) and colon epithelia (n=3 for each group of mice, so in total 6×3=18 RNA samples) by guanidinium thiocyanate-phenol-chloroform extraction and DNAse treatment, following the manufacturer specifications. Total RNA library preparation and sequencing were performed by Novogene Co., Ltd, as a lncRNA sequencing service, including lncRNA directional library preparation with rRNA depletion (Ribo-Zero Magnetic Kit), quantitation, pooling and PE 150 sequencing (30G raw data) on Illumina HiSeq 2500 platform. Filtering and trimming of the RNA seq data left around 230-300 million reads pairs /sample.

Gene expression analysis and differential splicing analysis was carried out by Novogene Co., Ltd, using the DESeq2 (v1.18.1) package^50^ and rMATS (v4.1.0)^51^ (parameters: --libType fr-firststrand −novelSS), respectively, on the mm10 mouse reference genome. For assessment cryptic splice sites, in house RNA-seq data and publicly available Rbm17 KO RNA-seq data (GEO GSE79020) were realigned on the mouse mm9 annotation file from Ensembl (version 67). The resulting BAM files were used to generate bigwig files, using bamCoverage (parameter: --normalizeUsing CPM) from Deeptools (v3.1.3)^52^ Coordinates of annotated and unannotated junctions were then retrieved from the BAM files using regtools (https://github.com/griffithlab/regtools). For each of the 6 conditions, unannotated junctions from the 3 replicates were pooled, before eliminating junctions present both in WT and KO conditions. Finally, junctions spanning over 20kb were filtered away as possible artefacts of the alignment. The remaining junctions, never shared between WT and mutant samples, were considered *de novo*. To quantify the strength of the splice sites, DNA sequences comprising the last 3 nucleotides of the exon and the first 6 nucleotides of the intron were recovered for the 5’ end of the junction. For the 3’ end of the junction, the sequences of the last 20 nucleotides of the intron and the first 3 nucleotides of the exon were recovered. Then, the splice sites strength was computed for each junction using the MaxEnt algorithm^53^. The MaxEnt algorithm was also applied to randomized junctions generated by applying the “shuffle” function of the bedtools suite (v.2.25.0)^54^ to the .bed files containing de novo junctions.

GSEA analysis was performed (with -nperm 1000 -permute gene_set -collapse false parameters) using the same expression matrix (v2.2.2), comparing ctrl and *Cbx3* KO samples at the crypt, villi and colon. Transcriptional signatures used for the analysis were extracted from the literature for lgr5 ISC signature^55^, enterocyte and Paneth cell signatures^56^ and GSEA data bases (MSigDB hallmark gene set).

### Fecal microbiota analysis by 16S rRNA gene sequencing

Genomic DNA was obtained from fecal samples using the QIAamp power fecal DNA kit (Zymo Research), and DNA quantity was determined using a TECAN Fluorometer (Qubit® dsDNA HS Assay Kit, Molecular Probes). The V3-V4 hypervariable region of the 16S rRNA gene was amplified by PCR using the following primers: a forward 43-nuclotide fusion primer 5’ **CTT TCC CTA CAC GAC GCT CTT CCG ATC T**AC GGR AGG CAG CAG3 consisting of the 28-nt illumina adapter (bold font) and the 14-nt broad range bacterial primer 343F and a reverse 47-nuclotide fusion 5’ **GGA GTT CAG ACG TGT GCT CTT CCG ATC T**TA CCA GGG TAT CTA ATC CT3’ consisting of the 28-nt illumina adapter (bold font) and the 19-nt broad range bacterial primer 784R. The PCR reactions were performed using 10 ng of DNA, 0.5 μM primers, 0.2 mM dNTP, and 0.5 U of the DNA-free Taq-polymerase, MolTaq 16S DNA Polymerase (Molzym). The amplifications were carried out using the following profile: 1 cycle at 94°C for 60 s, followed by 30 cycles at 94°C for 60 s, 65°C for 60 s, 72°C for 60 s, and finishing with a step at 72°C for 10 min. The PCR reactions were sent to the @Bridge platform (INRAe, Jouy-en-Josas) for sequencing using Illumina Miseq technology. Single multiplexing was performed using home-made 6 bp index, which were added to R784 during a second PCR with 12 cycles using forward primer (AATGATACGGCGACCACCGAGATCTACACTCTTTCCCTACACGAC) and reverse primer(CAAGCAGAAGACGGCATACGAGAT-index GTGACTGGAGTTCAGACGTGT). The resulting PCR products were purified and loaded onto the Illumina MiSeq cartridge according to the manufacturer instructions. The quality of the run was checked internally using PhiX, and then, sequences were assigned to its sample with the help of the previously integrated index. High quality filtered reads were further assembled and processed using FROGS pipeline (Find Rapidly OTU with Galaxy Solution) to obtain OTUs and their respective taxonomic assignment thanks to Galaxy instance (https://migale.inra.fr/galaxy). In each dataset, more than 97% of the paired-end sequences were assembled using at least a 10-bp overlap between the forward and reverse sequences. The following successive steps involved de-noising and clustering of the sequences into OTUs using SWARM, chimera removal using VSEARCh. Then, cluster abundances were filtered at 0.005%. One hundred percent of clusters were affiliated to OTU by using a silva138 16S reference database and the RDP (Ribosomal Database Project) classifier taxonomic assignment procedure. Richness and diversity indexes of bacterial community, as well as clustering and ordinations, were computed using the Phyloseq package (v 1.19.1) in RStudio software^57^. Divergence in community composition between samples was quantitatively assessed by calculating β-diversity index (UniFrac and weighted UniFrac distance matrices). For the heatmap, a negative binomial model was fit to each OTU, using DESeq2^50^ with default parameters, to estimate abundance log-fold changes (FCs). Values of *P* were corrected for multiple testing using the Benjamini-Hochberg procedure to control the false-discovery rate and significant OTUs were selected based on effect size (FC >|2|, adjusted *P* value <0.05).

### Mass spectrometry

#### Protein digestion

Immunoprecipitation eluates were digested following a FASP protocol^58^ slightly modified. Briefly, proteins were reduced using 100 mM DTT (dithiothreitol) for 1h at 60°C. Proteins were alkylated for 30 min by incubation in the dark at room temperature with 100 μL of 50 mM iodoacetamide. Samples were digested with 2μL of sequencing grade modified trypsin (Promega, WI, USA) for 16h at 37°C. Peptides were collected by centrifugation at 15,000 x g for 10 min followed by one wash with 50mM ammonium bicarbonate and vacuum dried.

#### NanoLC-MS/MS protein identification and quantification

Peptides were resuspended in 21 μL of 10% ACN, 0.1% TFA in HPLC-grade water prior MS analysis. For each run, 5 μL were injected in a nanoRSLC-Q Exactive PLUS (RSLC Ultimate 3000) (Thermo Scientific,Waltham MA, USA). Peptides were loaded onto a μ-precolumn (Acclaim PepMap 100 C18, cartridge, 300 μm i.d.×5 mm, 5 μm) (Thermo Scientific), and were separated on a 50 cm reversed-phase liquid chromatographic column (0.075 mm ID, Acclaim PepMap 100, C18, 2 μm) (Thermo Scientific). Chromatography solvents were (A) 0.1% formic acid in water, and (B) 80% acetonitrile, 0.08% formic acid. Peptides were eluted from the column with the following gradient 5% to 40% B (38 minutes), 40% to 80% (1 minute). At 39 minutes, the gradient stayed at 80% for 4 minutes and, at 43 minutes, it returned to 5% to re-equilibrate the column for 16 minutes before the next injection. Two blanks were run between each series to prevent sample carryover. Peptides eluting from the column were analyzed by data dependent MS/MS, using top-10 acquisition method. Peptides were fragmented using higher-energy collisional dissociation (HCD). Briefly, the instrument settings were as follows: resolution was set to 70,000 for MS scans and 17,500 for the data dependent MS/MS scans in order to increase speed. The MS AGC target was set to 3.10^6^ counts with maximum injection time set to 200 ms, while MS/MS AGC target was set to 1.10^5^ with maximum injection time set to 120 ms. The MS scan range was from 400 to 2000 m/z.

#### Data Analysis following nanoLC-MS/MS acquisition

Raw files corresponding to the proteins immunoprecipitated were analyzed using MaxQuant 1.5.5.1 software against the Human Uniprot KB/Swiss-Prot database 2016-01 ^59^. To search parent mass and fragment ions, we set a mass deviation of 3 ppm and 20 ppm respectively, no match between runs allowed. Carbamidomethylation (Cys) was set as fixed modification, whereas oxidation (Met) and N-term acetylation were set as variable modifications. The false discovery rates (FDRs) at the protein and peptide level were set to 1%. Scores were calculated using MaxQuant as described previously^59^. Peptides were quantified according to the MaxQuant MS1 signal intensities. Statistical and bioinformatic analysis, including volcano plot, were performed with Perseus software version 1.6.7.0 (freely available at www.perseus-framework.org). For statistical comparison, we set two groups, IP and negative control, each containing 3 biological replicates. We then retained only proteins that were quantified 3 times in at least one group. Next, the data were imputed to fill missing data points by creating a Gaussian distribution of random numbers with a standard deviation of 33% relative to the standard deviation of the measured values and 3 standard deviation downshift of the mean to simulate the distribution of low signal values. We performed a T-test, and represented the data on a volcano plot (FDR<0.05, S0=1).

### SDS-PAGE and immunoblotting

Intestinal epithelial cells purified from mice (at least n = 3 separate experiments) were lysed at 4 °C in a buffer containing 25mM Tris pH 7.5, 1mM EDTA, 0.1mM EGTA, 5mM MgCl2, 1% NP-40, 10% Glycerol, 150mM NaCl, and then cleared by centrifugation at 14,000 rpm for 30 min at 4 °C. Proteins were separated on SDS–PAGE gels and transferred to nitrocellulose membranes by standard procedures. Mouse anti-progerin monoclonal antibody (13A4DA, sc-81611, Santa Cruz), anti-HP1 *α* (2H4E9, Novus Biologicals), HP1 *β* (1MOD-1A9, Thermo Scientific) and HP1 *γ* (2MOD-IG6, Thermo Scientific) and *γ* Tubulin (4D11, Thermo Scientific) antibodies.

### Statistical Analyses

Statistical analyses were performed with the GraphPad Prism soft-ware. Differences between groups were assayed using a two-tailed Student’s t-test using GraphPad Prism. In all cases, the experimental data were assumed to fulfill t-test requirements (normal distribution and similar variance); in those cases, where the assumption of the t-test was not valid, a nonparametric statistical method was used (Mann–Whitney test). The differences between three or more groups were tested by one-way ANOVA. A p-value less than 0.05 was considered as significant. Error bars indicate the standard error of the mean.

## Data access

RNA-seq data, the derived mm9.bigwig files, and the *de novo* junction.bed files generated in this study have been submitted to the NCBI Gene Expression Omnibus (GEO; http://www.ncbi.nlm.nih.gov/geo/) under accession numbers GSE192800. **Access will be granted to reviewers with the following token: mdkrywoyphyplwh**

## ACKNOWLEDGMENTS

We thank Florence Cammas for providing the *Cbx3*^*fl*/fl^ mice. We thanks Dr Zhou Zhongjun (Department of Biochemistry, Li Ka Shing Faculty of Medicine The university of Hong Kong) for providing us MEF from HGPS mice. We thank Dr Cohen-Tannoudji for providing the Villin-CreERT2 mice (Institut Pasteur, Paris, France)

